# Early spatiotemporal evolution of the immune response elicited by adenovirus serotype 26 vector vaccination in mice

**DOI:** 10.1101/2024.10.18.618988

**Authors:** Eryn Blass, Alessandro Colarusso, Malika Aid, Rafael A. Larocca, R. Keith Reeves, Dan H. Barouch

**Author notes:** Department of Medical Oncology, Dana-Farber Cancer Institute, Harvard Medical School, Boston, Massachusetts, USA. Department of Surgery, Duke University School of Medicine, Durham, North Carolina, USA. Center for Human Systems Immunology, Duke University, Durham, North Carolina, USA.

## Abstract

As the first responder to immunological challenges, the innate immune system shapes and regulates the ensuing adaptive immune response. Many clinical studies evaluating the role of innate immunity in initiating vaccine-elicited adaptive immune responses have largely been confined to blood due to inherent difficulty in acquiring tissue samples. However, the absence of vaccine-site and draining lymph node information limits understanding of early events induced by vaccination that could potentially shape vaccine-elicited immunity. We therefore utilized a mouse model to investigate the spatiotemporal evolution of the immune response within the first 24 hours following intramuscular adenovirus serotype 26 (Ad26) vector vaccination in tissues. We show that the Ad26 vaccine-elicited innate immune response commences by one hour and rapidly evolves in tissues and blood within the first 24 hours as reflected by the detection of cytokines, chemokines, cellular responses, and transcriptomic pathways. Furthermore, serum levels of IL-6, MIG, MIP-1α, and MIP-1β at 6 hours post-vaccination correlated with the frequency of vaccine-elicited memory CD8^+^ T cell responses evaluated at 60 days post-vaccination in blood and tissues. Taken together, our data suggests that the immune response to Ad26 vector vaccination commences quickly in tissues by one hour and that events by as early as 6 hours post-vaccination can shape vaccine-elicited CD8^+^ T cell responses at later memory time points.

**IMPORTANCE:** Prior studies have largely concentrated on innate immune activation in peripheral blood following vaccination. In this study, we report the detailed spatial and temporal innate immune activation in tissues following Ad26 vaccination in mice. We observed rapid innate activation rapidly not only in peripheral blood but also in draining lymph nodes and at the site of inoculation. Our findings provide a more detailed picture of host response to vaccination than previously reported.

## INTRODUCTION

Innate immunity plays a critical role as an initial barrier to infection and forms an integral component in the initiation and development of adaptive immune responses. As the generation of protective adaptive immune responses is critical to the development of successful vaccines, understanding the bridge between innate and adaptive immunity provides greater insights into the immunological mechanisms of vaccination and how immune responses are ultimately tailored. Adenovirus (Ad) vectors have been extensively studied for vaccine development for infectious diseases such as HIV^1^, Zika^2^, Ebola^3^, and SARS-COV2 ^4,5^. CD8^+^ T cell responses are strongly induced by Ad vectors and as such this vaccine platform has the utility of being used for T cell-based vaccines. Inducing CD8^+^ T cell responses to conserved T cell epitopes has the ability to provide cross-protective immunity to evolving pathogens which would otherwise escape neutralizing antibodies, such as SARS-COV2 ^6,7^. Secondly, the ability to induce robust anti-tumor CD8^+^ T cells also positions their application in therapeutic cancer vaccines as demonstrated in mouse studies^8,9^ and recent phase I clinical trials^10,11^.

Prior studies have evaluated how innate immune induction coordinates with vaccine-elicited adaptive immune responses, however many have been restricted to the study of peripheral blood in humans ^12–15^, with limited investigations in tissues in mice ^16–20^. These studies have not addressed the earliest kinetics across tissues. We therefore sought to elucidate the early spatiotemporal evolution of the immunological response following Ad vector vaccination. We aimed to integrate the early immune response with the induction of CD8^+^ T cell responses to understand underlying factors that influence immunogenicity of T cell-based vaccines.

We found that the initial wave of the immune response following intramuscular Ad26 vaccination commences by one hour and develops quickly over the first 24 hours across tissues and blood. Serum cytokines at six hours correlated with the frequency of vaccine-elicited CD8^+^ T cells at 60 days post-vaccination, suggesting that immunological events within the first few hours already have the potential to shape memory CD8^+^ T cell formation. These studies lay foundation for more detailed mechanistic studies into vaccine-elicited innate immunity and its integration with ensuing adaptive immune responses.

## RESULTS

### The initial wave of immune response following Ad26 vector vaccination in mice occurs within the first 24 hours

We first aimed to understand the general kinetics of the early immunological response following viral vector vaccination. C57BL/6 mice were vaccinated intramuscularly in with a prototype Ad26 vaccine vector expressing SIVgag (Ad26-SIVgag), and we conducted a time course study focusing on the first 24 hours post-vaccination (**Figure 1A**) at 1, 3, 6, 12, 24 hours and an additional time point later at 72 hours to reflect the likely waning innate immune response. As cytokines and chemokines are critical for the initiation, coordination, and resolution of inflammation, we assessed the induction kinetics of cytokines and chemokines via multiplex bead-based ELISA (Luminex) assays in serum post-vaccination.

**Figure 1.**
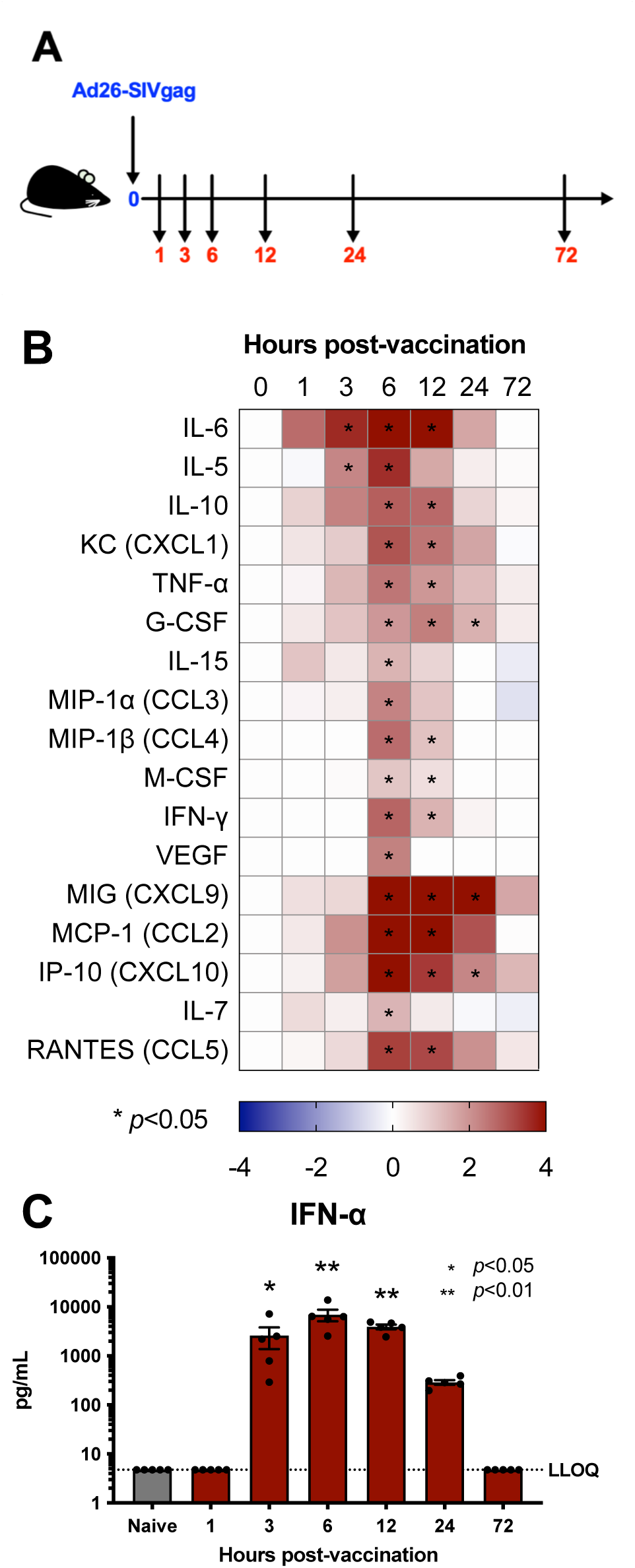
Intramuscular adenovirus serotype 26 vector vaccination rapidly induces serum cytokines, chemokines, and interferon. C57BL/6 mice were immunized intramuscularly with 1×10^10^vp of Ad26-SIVgag. A) Study outline, B) Cytokines and chemokines detected in serum via Luminex are shown as a heat map of log2 fold change (LOG_2_FC) of the group average over the average naive reading, C) IFN-α levels as measured via IFN-α ELISA. N of 5 per group. Kruskall-Wallis test with Dunn’s corrections for multiple comparisons.

We found that the cytokine response initiates by significant IL-6 detection at 3 hours (*p*=<0.05) (**Figure 1B).** Peak responses occurred around the 6 hour time point (IL-6, IL-5, IL-10, CXCL1, TNF-α, MIP-1α, MIP-1β, MIG, MCP-1, IP-10, IL-7, RANTES, all at least *p*=<0.05) and a later set exhibiting a more prolonged detection at 24 hours (G-CSF, MIG, IP-10, all at least *p*<0.05). Similar kinetics were observed for IFN-α in serum (3-12 hours, all at least *p*=<0.05) (**Figure 1C**). Overall, responses began to wane by 24 hours and were mostly close to baseline by 72 hours (**Figures 1B and 1C**). These data suggest that the initial cytokine and chemokine induction occurs in a rapid and transient fashion following intramuscular Ad26 vaccination, thus quickly coordinating the initiation and integration of immune responses.

### Ad26 vector vaccination results in rapid evolution of multiple immunologic pathways across blood and tissues within the first 24 hours post-vaccination

We then sought to garner a global picture of how the immunological response to Ad26 vector vaccination develops across time and space in more detail. We collected blood and tissues to survey immune response kinetics by bulk RNA-seq transcriptomic profiling, including the site of vaccination in muscle, the draining iliac lymph node (dLN) for priming of adaptive immune responses, and additionally peripheral blood (**Figure 2A)**. As cell-to-cell signaling is critical for initiating the coordination of immune responses, we first evaluated gene expression levels of cytokines and chemokines across all time points and compartments.

**Figure 2.**
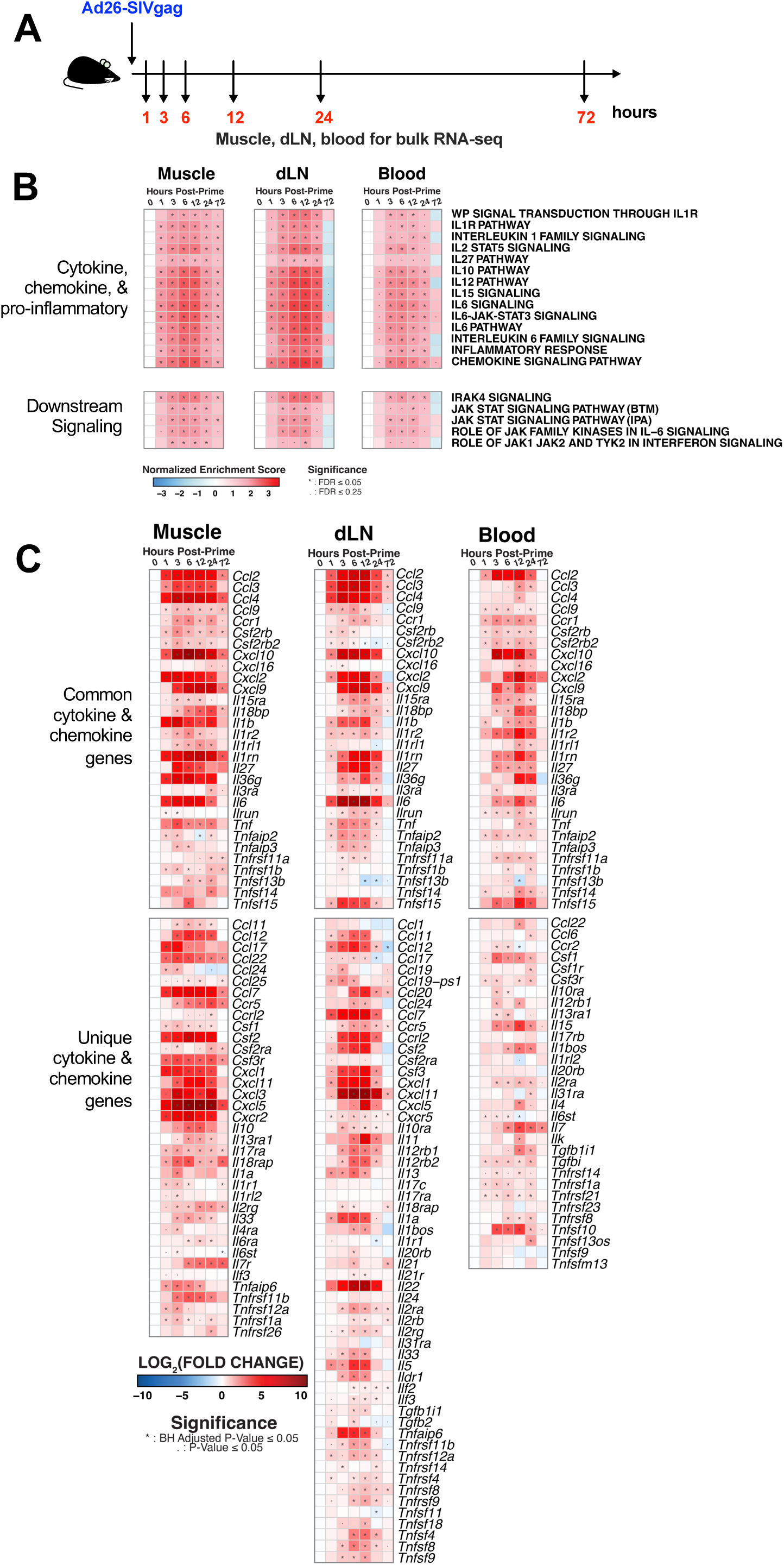
Ad26 vector vaccination broadly stimulates immunologic signaling pathways across tissues within the first few hours following vaccination. C57BL/6 mice were immunized intramuscularly with 1×10^10^vp of Ad26-SIVgag and samples were harvested for immune analysis via bulk RNA-seq across muscle, dLN, and blood. A) Gene Set Enrichment Analysis (GSEA) analysis of immune signaling pathways as measured by Normalized Enrichment Score (NES) from naive, B) individual genes related to cytokines and chemokines (LOG_2_FC from naive). N of 5 per group.

Across all three compartments we observed rapid upregulation of a wide variety of pathways for cytokine and chemokine signaling, and related downstream signaling **(Figure 2B, top and bottom panels, respectively)**. These included multiple pathways for proinflammatory IL1 and IL6 signaling, in addition to IL12, IL-15 and chemokine signaling (**Figure 2B top panel, Supplementary Tables 1-3)**. At 1 hour we observe d significant enrichment of these pathways in muscle and the dLN compared to blood. These pathways were largely downregulated in the dLN and blood by 72 hours **(Supplementary Table 2)**, while still persisting in the muscle (IL-12 pathway *NES*=1.73 *p*<0.0001, *FDR*<0.001; IL-15 pathway *NES*=1.67, *p*<0.0001, *FDR*<0.01; IL-6 signaling *NES*=1.73 *p*<0.0001, *FDR*<0.001) **(Supplementary Table 1)**. These data highlight the rapid induction of immune responses at the vaccine site and dLN, with following detectable responses in blood hours later.

We then evaluated individual cytokine and chemokine gene expression. We observed a number of significantly upregulated genes were shared across all three compartments with a general peak around 3-12 hours: *Ccl2, Ccl3, Ccl4, Ccl9, Cxcl10*, *Cxcl9*, *Il1b*, *Il6*, and *Tnf* (**Figure 2C top panel, Supplementary Tables 4-6**). While some degree of commonality existed across compartments, we observed clear differences in tissue-specific gene expression (**Figure 2C bottom panel**). In muscle pro-inflammatory *Cxcl3* (3hrs *p*<0.0001, 24hrs peak *p*<0.0001) and anti-inflammatory *Ccl17* (1hr *p*<0.01, 3hrs *p*<0.001), *Ccl22* (1hr *p*<0.001, 3hrs *p*<0.0001, 6hrs *p*<0.0001), and *Il10* (6hrs *p*<0.01, 12hrs *p*<0.001) were upregulated early, suggesting a balancing of induced immune responses.^21^ Further, while overall response waned by 72 hours, the muscle still exhibited prolonged expression of *Ccl2* (*p*<0.0001), *Ccl4* (*p*<0.01), *Cxcl10* (*p*<0.001), *Cxcl9* (*p*<0.01), *Ccl22* (*p*<0.05), *Ccl7* (*p*<0.0001), and *Cxcl5* (*p*<0.05) **(Figure 2C, Supplemental Table 4).**

In the dLN, unique cytokine and chemokines related to lymphocyte response and trafficking were marked by upregulation of *Ccl19* (3hrs *p*<0.0001), *Ccl20* (6hrs *p*<0.0001, 12hrs peak *p*<0.0001), *Il11* (6 hrs *p*<0.01, 12hrs peak *p*<0.0001), *Il13* (1hr *p*<0.01, 6hrs peak *p*<0.01), *Il22* (3hrs *p*<0.0001, 6-12hrs peak *p*<0.0001), and *Il5* (1hr *p*<0.05, 6-12hr peak *p*<0.0001) **(Figure 2C bottom panel, Supplemental Table 5)**. While in blood, gene expression related to immune proliferation was uniquely elevated as reflected by *Il15* (3hrs *p*<0.0001, 12hrs peak *p*<0.0002) and *Il7* (6hrs *p*<0.0001, 12hrs peak *p*<0.0001). Further, others trended towards a later peak response in blood compared to other compartments such as *Cxcl2* (12 hrs p<0.0001) and *Tnf* (12 hrs p<0.0001, 24 hrs p<0.01), and *Ccl22* at 12hrs (*p*<0.05) in blood versus 1 hr (*p*<0.0001) in muscle **(Figure 2B bottom panel, Supplemental Table 6).**

When we considered integrative immune processes across compartments. We first observed more common chemokine genes upregulated between in the muscle and dLN **(Figure 2C bottom panel)**. *Ccl7, Ccl12, Cxcl1, Cxcl11, and Cxcl5* are chemoattractants for lymphocytes and monocytes, together suggesting the initiation of monocyte trafficking and differentiation within hours in these two compartments. *Csf1* was upregulated between muscle and blood, which has a role in stimulating proliferation and differentiation of macrophages **(Figure 2C bottom panel)**. However, blood overall had fewer overlapping genes with either the muscle or dLN compartments, The overlap in gene expression pattern between muscle and draining lymph node could be reflective of the rapid spread of immune response and coordination between the injection site and its draining lymph node.

Following our analysis of cytokine and chemokine signaling, we evaluated the enrichment of interferon family genes as type I interferon signaling has been shown to shape the development of T cell magnitude and polyfunctionality following vaccination with some Ad vector serotypes^22^. We observed significant enrichment of many pathways associated with interferon responses and signaling across all compartments (**Figure 3A, Supplemental Tables 1-3**). Key downstream signaling genes *Irf9 (p<0.0001)* and *Isg15 (p<0.0001)* were upregulated by 3 hours in the muscle, dLN, and blood, alongside other associated genes. Initiating the response, production of IFN-α occurred only in the dLN as transcripts for multiple IFN-α subtypes were significantly induced starting at 1 hour: *Ifna1, Ifna2, Ifna4, Ifna5, Ifna6, Ifna7, Ifna9, Ifna11, Ifna12, Ifna13, Ifna14,* and *Ifna15* (all at least *p*<0.05) (**Figure 3B bottom panel, Supplementary Tables 3-6**). Our data supports and extends prior findings ^12,16,23,24^ by showing that rapid production of IFN-α in the draining lymph node by one hour likely results in systemic upregulation of interferon pathways by 3 hours post-vaccination.

**Figure 3.**
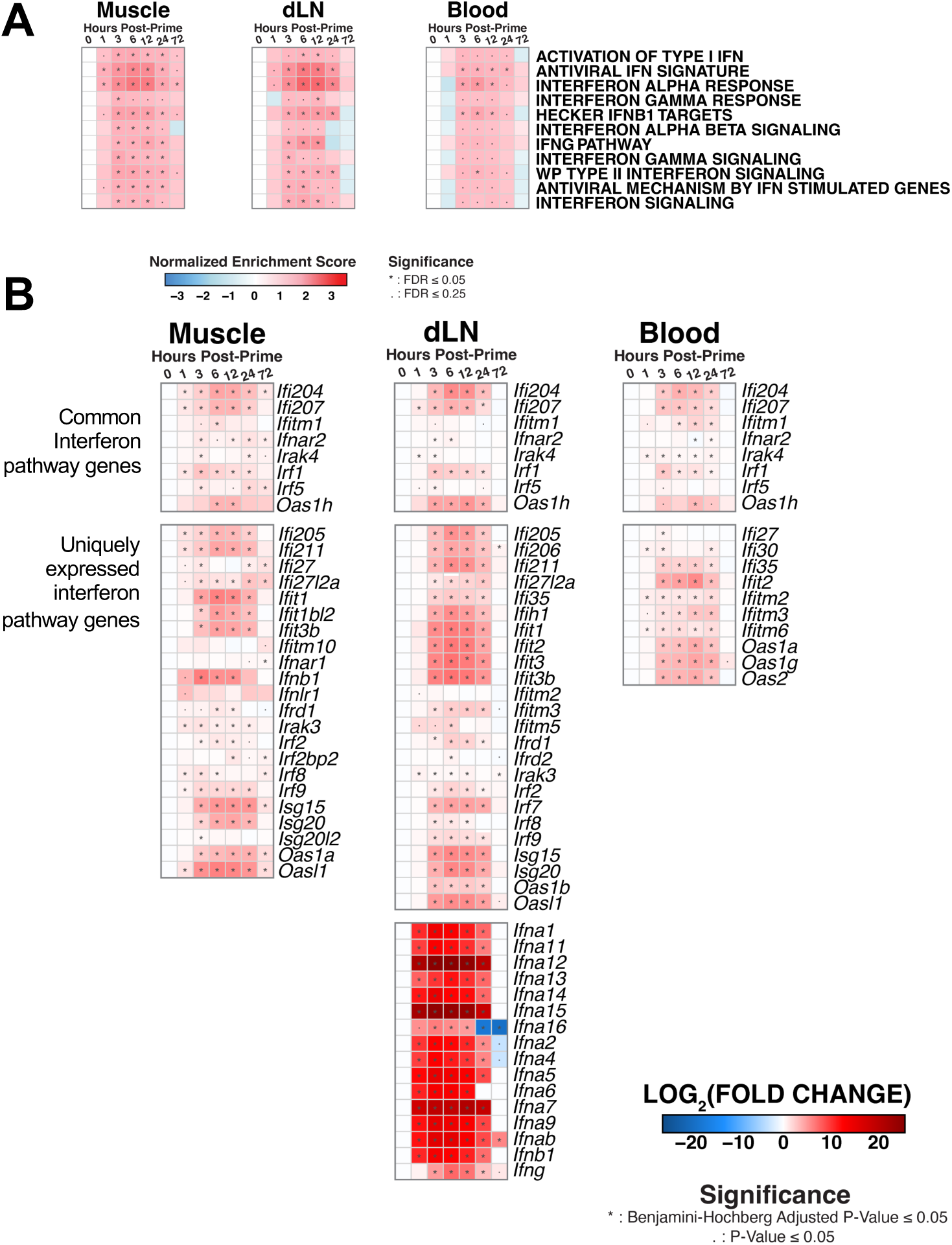
Interferon pathways are rapidly upregulated across tissues. Interferon responses across tissues were assessed by A) GSEA of interferon signaling pathways measured by NES as compared to naive and B) individual genes in interferon signaling module as measured by LOG_2_FC from naive. N of 5 per group.

Together these early pathway data suggest that immunological signaling pathways are significantly enriched not only at the injection site but also spreading to the draining lymph node as early as one hour post-Ad26 intramuscular vaccination, highlighting the rapid coordination of vaccine-induced immune responses. Furthermore, the patterns of overlap between muscle and dLN, but to a lesser extent blood, suggests that blood alone may not fully capture the extent of immunological responses.

### Myeloid cells are early responders to Ad26 intramuscular vaccination

Integrating our signaling data with cellular responses, we next sought to understand the immune cell components that could be initially driving and responding to the cytokine and chemokine signals by evaluating pathways for immune cell populations across all three compartments. M1 macrophage signatures were most consistently enriched post-vaccination with initial detection by 1 hour post-vaccination in muscle (*NES*=1.61, *FDR*<0.05) following by dLN at 3hrs (*NES*=1.99, *FDR*<0.001) and blood at 6 hrs (*NES*=1.66, *FDR*<0.05) (**Figure 4A**). We also observed rapid enrichment of the Activated Dendritic Cell signature pathway in muscle by 1 hour (*NES*=1.71, *FDR*<0.05), then followed by 3 hrs in dLN (*NES*=2.02, *FDR*<0.001) and blood (*NES*=1.69, *FDR*<0.05) **(Figure 4A)**.

**Figure 4.**
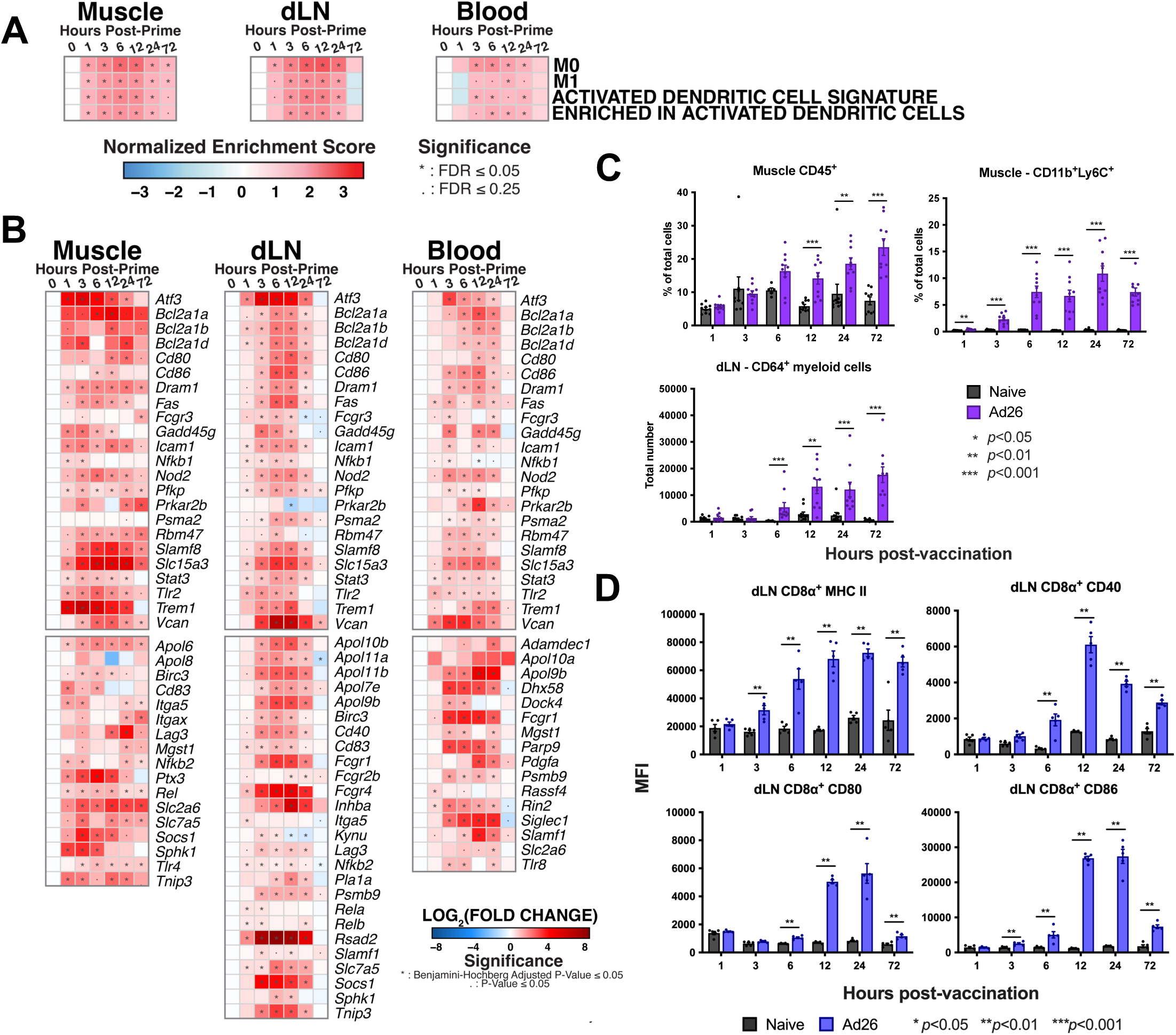
Early immunological responses are driven by myeloid cells. Bulk RNA-seq data was assessed for A) immune cell signatures by GSEA, and B) related individual genes. To evaluate cellular responses via flow cytometry, C57BL/6 mice were immunized intramuscularly with 1×10^10^ vp of Ad26-SIVgag. Muscle, draining lymph node, and blood, were collected and immunological responses were profiled by C) Total frequency of CD45^+^ cells in muscle, D) Total frequency of CD11b^+^Ly6C^+^ myeloid cells in muscle, E) Total number of CD11b^+^Ly6C^+^CD64^+^ monocyte-derived dendritic cells in dLN. F) Surface expression of MHC II (I-A/I-E), G) CD80, H) CD86, and I) CD40 on dLN CD8α^+^ dendritic cells, measured as Median Fluorescence Intensity (MFI). N of 5-10 per group, Mann-Whitney *U*-test.

Within the dLN at 6 hours we observed continued enrichment of the Activated Dendritic Cell pathway (*NES*=2.26, *FDR*<0.0001) with leading edge genes *Irf7, Cd40, Cd80, Cd86, Ccl19, Cxcl10, Cxcl11*, and the Enriched in Activated Dendritic cells (NES=2.08, *FDR*<0.001) pathway including genes *Il18, Il1b* **(Figure 4A**, **Figure 4B, Supplemental Table 5)**. These pathways suggest DC activation and maturation commencing within the dLN by 6 hours post-vaccination.

In order to confirm the transcriptomic changes that were observed post-vaccination, we profiled the response kinetics of myeloid cell populations. While the total frequency of CD45^+^ cells was not significantly higher at one hour post-vaccination in muscle **(Figure 4C)**, we observed a significant increase in the frequency of CD11b^+^Ly6C^+^ immune cells in at 1 hour post-vaccination (*p*<0.01) continuing through to 72 hours (**Figure 4C**), suggesting an accumulation of inflammatory monocytes. While trends emerged earlier, starting at 12 hours post-vaccination we observed a progressive significant increase in CD45^+^ cells in the muscle (*p*<0.001), reflecting increased immune cell recruitment to the initial injection (**Figure 4C**). Together these data indicate that while total frequency of immune cells many not change significantly in the initial hours post-vaccination, changes occur within its composition. Of these immune cells, inflammatory monocytes are among the earliest responders at the vaccination site, followed by immune cell recruitment to the vaccine site.

We then considered the kinetics of immunologic responses in the dLN. We observed the appearance of a CD11b^+^Ly6C^+^CD64^+^ population in dLN starting by 6 hours post-vaccination (*p*<0.001) (**Figure 4C**). CD64 can be expressed on monocytes, macrophages, and monocyte-derived dendritic cells (mo-DC)^25^. Previously published data has shown that antigen-carrying mo-DC were also found in the dLN 24 hours following subcutaneous vaccination with other Ad vector serotypes.^16^ The appearance of this population in the dLN occurred following detection of CD11b^+^Ly6C^+^ inflammatory monocytes in muscle. Our data support prior findings and by extension suggest that Ly6C^+^ inflammatory monocytes and CD64^+^ myeloid cells may play a prominent and early role in the first few hours following intramuscular Ad26 vector vaccination.

It is known that dendritic cell cross-presentation of antigen is critical for the induction of CD8^+^ T cell responses following Ad vector vaccination, which includes the lymph node resident CD8α^+^ DC population.^16,26^ We therefore evaluated the response kinetics of the CD8α^+^ DC subset (CD11c^+^CD8^+^B220^-^). We observed significantly increased expression of MHC II (I-A/I-E) first at 3 hours (*p*<0.01) **(Figure 4D)** and CD86 (*p*<0.01), followed by co-stimulatory markers CD40 (*p*=0.01), CD80 (*p*=0.01) at 6 hours post-vaccination, with reduced expression by 72 hours. Taken together, our studies showed rapid trafficking of myeloid cells into the muscle injection site within hours following vaccination. Considering the vast array of cytokines and chemokines that can be released by monocyte and macrophage populations, they may play a substantial role in promoting Ad26 the initial vaccine-elicited immune responses in the first few hours following intramuscular vaccination.

### CD8^+^ T cell immunogenicity can be shaped by 6 hours following Ad26 vaccination

As innate immunity can shape and regulate adaptive immune responses, we sought to understand how markers of early immune responses could serve as an indicator of vaccine-elicited CD8^+^ T cell responses. We vaccinated mice with Ad26-SIVgag and collected serum for protein-level cytokine analysis (Luminex) at 6 hours post-vaccination. We chose this time point as we previously observed the broadest degree of cytokine and chemokines detection in serum. We then evaluated the induction of SIVgag-specific CD8^+^ T cell responses via tetramer binding assays for the immunodominant SIVgag H-2D^b^ epitope, AL11^27^ at 60 days post-vaccination in blood and tissues (**Figure 5A**). We found that the frequency of AL11-specific CD8^+^ T cells at 60 days post-vaccination in blood, dLN, and spleen positively correlated with the levels of IL-6 (*p*=0.0101, *p*=0.0011, *p*=0.0033, respectively), MIG/CXCL9 (*p*=0.0212, *p*=0.0229, *p*=0.0138), MIP-1α (*p*=0.0087, *p*=0.0040, *p*=0.0002), and MIP-1β (*p*=0.0089, *p*=0.0227, *p*=0.0081) at 6 hours post-vaccination (**Figure 5B**). Together these data suggest that immunological events occurring by 6 hours post-vaccination already have the ability to shape the Ad26-vaccine elicited CD8^+^ T cell response, including the generation of memory T cell responses.

**Figure 5.**
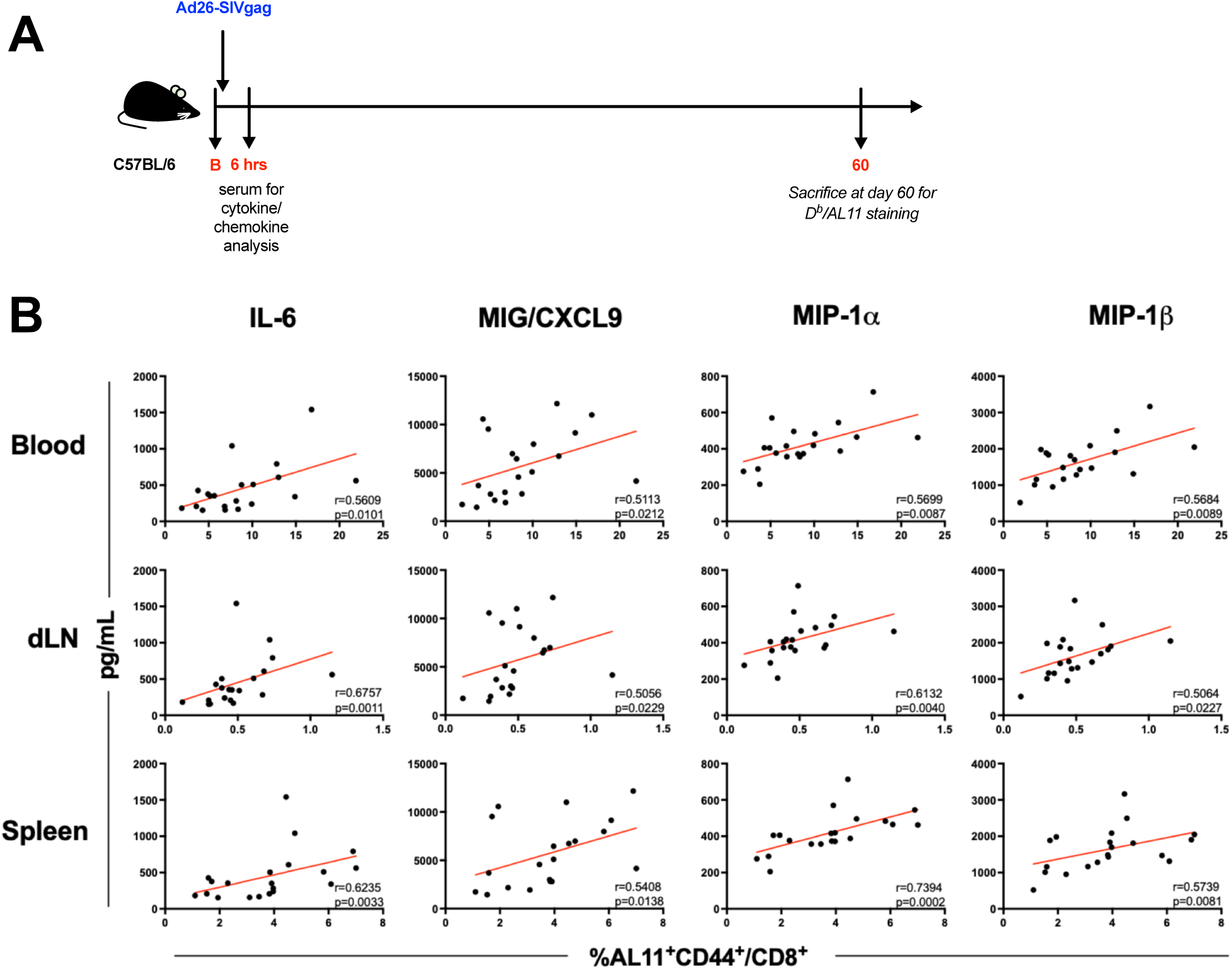
Serum cytokines at 6 hours post-vaccination are predictive of vaccine-elicited CD8^+^ T cell responses. C57BL/6 mice were immunized with 1×10^10^ vp of Ad26-SIVgag. Cytokine and chemokine protein levels were evaluated in serum via Luminex at 6 hours post-vaccination. A) Outline of study. B) Correlation of serum cytokine levels measured via Luminex with frequency of SIVgag-specific CD8^+^ T cell responses measured via H-2D^b^ AL11 tetramer binding assays in blood, dLN, and spleen at 60 days post-vaccination. N of 20 per group. Spearman rank test.

## DISCUSSION

Adenovirus vectors have demonstrated their utility as vaccine platforms due to their ability to stimulate robust immune responses following vaccination. While much is known about immune responses elicited by Ad vectors, open questions remain as to what immediate immunological events occur following vaccination and in particular how these early events are unfolding in tissues that could potentially shape and regulate vaccine-elicited immunity.

Prior studies have investigated the rapidity of immune responses elicited by vaccination ^16,18–20,28,29^. We hypothesized that we would observe immune responses unfold within hours following Ad26 vaccination, initiating from the site of vaccination, to the draining lymph node, and systemically reflected in blood. Building upon prior observations demonstrating Ad26-induced serum cytokine responses at 24 hours post-vaccination in non-human primates^23^ and humans^30^ we now use a mouse model to detail the evolution of the earliest immune responses during the first 24 hours post-vaccination across blood and tissues.

By transcriptomic analysis, enrichment of TNF, IL1, and IL6 pro-inflammatory pathways by one-hour post-vaccination across tissues suggests a systemic rapid coordination of immune response. These pro-inflammatory pathways likely initiate the cascade of increased cytokine and chemokine gene expression and protein levels at 3-12 hours. These responses typically peaked within the first 24 hours, indicating that while some degree of innate immune responses can be detected a day post-vaccination, earlier timepoints may be of more interest for surveying a greater breadth and magnitude of innate immune responses. Furthermore, the breadth of induced immune signaling pathways may suggest that Ad26 can broadly stimulate the induction of immune responses, which may contribute to its potent immunogenicity.

Some pathways exhibited commonality across anatomic compartments, suggesting key unifying immunological events. However, we also observed tissue-specific features. Innate immune responses waned quickly in blood but persisted in muscle, likely due to ongoing immune recruitment as suggested by continued enrichment of myeloid cell gene signatures and detectable CD11b^+^LyC^+^ cells in muscle at 72 hours. Additionally, overlap is observed more between muscle and dLN in comparison to blood suggesting that immune responses may be tightly coordinated between these two compartments. This also suggests that sampling of blood for the study of innate immune responses, while showing some unified immune responses, may not entirely reflect key immunological events that determine vaccine immunogenicity that are uniquely tissue-located.

When we evaluated potential serum biomarkers of vaccine immunogenicity, we observed that serum levels of IL-6, MIG/CXCL9, MIP-1α, and MIP-1β at 6 hours correlated with the frequency of SIV-gag AL11-specific CD8^+^ T cell responses in blood and tissues at 60 days post-vaccination. In hand, we observed *Il6*, *Cxcl9*, *Ccl3*, and *Ccl4* gene expression in all three compartments, but more strongly upregulated in Muscle and dLN. IL-6 is a pleiotropic cytokine that has a role in various facets of pro-inflammatory immune responses and immune coordination. IL-6 has been shown to be produced rapidly by macrophages and dendritic cells following systemic intravenous administration of Ad5^31^. IL-6 has been shown to play a role in promoting Ad5-induced CD8^+^ T cell responses following co-administration with HDAC inhibitors ^8^. Our data show a potential role of IL-6 in the coordination of Ad26 vaccine-elicited CD8^+^ T cell responses, which is likely multifactorial. Although the mechanism was not defined, MIP-1α (CCL3) has been shown to increase CD4^+^ T cell responses when encoded alongside vaccine antigens in an Ad5 vaccination mouse model ^32^. While we were able to identify serum biomarkers of vaccine immunogenicity, deeper mechanistic studies are warranted to understand the role of these cytokine and chemokine pathways in shaping Ad26 vaccine-elicited CD8^+^ T cell responses.

While we focused on Ad26, a prior study in mice evaluated transcriptomic responses at 8, 24, and 72 hours post-vaccination in the draining lymph node following subcutaneous vaccination with a variety of Ad vector serotypes including Ad5, Ad28, Ad35, chAd3, chAd63, sAd11, sAd16^16^. In line with that study, we observe similar response kinetics in our data post-intramuscular Ad26 vaccination in the dLN. Another study analyzed early immune responses in muscle and dLN following ChAd155 intramuscular vaccination in mice in which cytokine responses were detected at 1 hour in muscle ^20^. This observation is concordant with findings with Ad26, however we extend this knowledge by evaluating the broader immunologic transcriptomic networks involved in the immune cascade, and integration of serum biomarkers with vaccine immunogenicity.

In our study we used a mouse model due to the ease of tissue sampling to investigate the kinetics of tissue-specific immunity. Unlike its widely distributed expression in humans, CD46 expression in mice is limited to testes and retinal tissue. CD46 is a primary entry receptor for Ad26^33^. Prior studies investigating differences in T cell phenotypes with CD46 utilizing vectors have shown similarities between T cell responses in C57BL/6 and CD46 transgenic mice engineered to express the CD46 receptor, suggesting that the lack of CD46 does not dramatically impact the induction or shaping of vaccine-induced T cell responses in this mouse model^34^. Intramuscular injection of Ad26 induces transgene expression in the muscle^35^. These data suggest that regardless of the absence of CD46 expression, Ad26 can enter cells at the site of injection in the muscle and as such potentially may use alternative entry mechanisms in this vaccination route in mice.

Our study design used bulk RNA-seq to survey a large number of samples across tissues and timepoints. A limitation of this approach is that bulk RNA-seq cannot capture immune cell heterogeneity and specific functional assignment on a per-cell basis. Myeloid cell populations can differentiate into a variety of states and subsets in the context of inflammation, thus minor and novel subsets cannot be defined through this approach. Moving forward, studies utilizing single-cell approaches will provide greater depth of immunological cell states and their corresponding functional signatures.

Taken together, our data show that the innate immune response elicited by Ad26 vaccination commences by one hour post-vaccination and rapidly evolves within the first 24 hours across the site of vaccination, the draining lymph node, and blood. Immunologic pathways suggest rapid coordination of immune responses, immune cell trafficking, and cellular responses. While CD8α^+^ cross-presenting DCs are critical for the induction of CD8^+^ T cell immunity, the monocyte/macrophage lineage may be a significant contributor to initiating immune responses following intramuscular Ad26 vector vaccination. Furthermore, immunological events occurring within a few hours post-vaccination shape the vaccine-elicited memory CD8^+^ T cell response. These data highlight the rapidity of the innate immune system in tissues in initiating and shaping the ensuing vaccine-elicited adaptive immune response and merits deeper investigation into early mechanisms of immune induction for understanding rational vaccine design. Future studies should also define the early spatiotemporal evaluation of innate and adaptive immune responses with other vaccine platforms.

## AUTHOR CONTRIBUTIONS

E.B. and D.H.B. designed the studies. E.B. conducted all animal studies, immunologic studies and corresponding analyses. A.C. and M.A. performed the computational analyses. R.A.L. provided experimental assistance and guidance. R.K.R. contributed to study conception and assisted with data interpretation. E.B. and D.H.B. wrote the paper with all co-authors.

## ACKNOWLEDGEMENTS

We would like to thank Peter Abbink, Rebecca Peterson, Noe Mercado, Abishek Chandrashekar, Justin Iampietro, and Zi Han Kang for advice and technical assistance. We thank the NIH Tetramer Core Facility for provision of AL11 monomers, Zach Herbert and the Dana-Farber Molecular Biology Core Facility for assistance and advice with RNA-seq experiments, and Michelle Lifton and Rachel Hindin of the Center for Virology and Vaccine Research flow cytometry core facility. We acknowledge support from NIH grants AI128751, AI149670, AI164556, AI169615, AI177687.

## MATERIALS AND METHODS

### Immunizations

Female C57BL/6 mice were purchased from Jackson Laboratories (Bar Harbor, ME). Replication-incompetent, recombinant E1/E3-deleted adenovirus serotype 26 (Ad26) vectors were previously constructed ^34,36^. Mice were immunized by bilateral intramuscular injection into the hind leg quadriceps with 10^10^ viral particles (vp) per mouse. All experiments were performed with approval from the BIDMC Institutional Animal Care and Use Committee (IACUC).

We performed a dose-titration experiment to determine the optimal dose for investigating innate immune responses. C57BL/6 mice were vaccinated intramuscularly into the hind leg quadriceps with escalating doses of an Ad26 vector expressing SIVgag: 1×10^8^, 1×10^9^, 1×10^10^ viral particles (vp). We observed that at 8 hours post-vaccination some cytokines were below the limit of detection at a dose lower than 1×10^10^ vp (*unpublished*). This suggests immune responses may be low and potentially below assay detection limits depending on vector dose. Therefore for these studies we used a 1×10^10^ vp dose for evaluating innate immune responses in order to better detect low-level immune responses.

### Transcriptomic analyses

Tissue samples were collected into RNAlater (Invitrogen). Tissue samples were then transferred to QIAzol and homogenized with a TissueLyzer using 5mm steel beads (all Qiagen). Blood was processed as outlined in the sample collection section, with cell pellets resuspended in QIAzol. Total RNA was extracted according to the QIAcube HT RNA extraction protocol (Qiagen). The Dana-Farber Molecular Biology Core Facility evaluated RNA quality via the Agilent 2100 Bioanalyzer (Agilent Technologies) and prepared RNA-seq libraries. Single-end 75bp libraries were barcoded for multiplexing and sequenced with 20,000 reads per sample on an Illumina NextSeq 500.

#### RNA-seq analysis

All operations were performed on locally, in R (version 4.3.1) ^37^. To slightly reduce noise, raw RNA counts were filtered such that genes with counts greater than 0 across all animals were preserved for further analysis. Initial principal component analysis (PCA) and data visualization was performed on normalized counts, using the “plotPCA()” function in R’s DESeq2 (version 1.40.2) ^38^. Due to the robust clustering observed in the PCA, where the first principal component (PC1) explained approximately 80% of the variance, each tissue was analyzed separately. This approach allowed us to focus on tissue-specific responses without being confounded by inter-tissue variability. Counts were then normalized using the “deseq()” function. Differential gene expression was computed for post-vaccination timepoints by contrasting each timepoint to baseline/pre-vaccination with the “results()” function in DESeq2 ^38^. Parameters in DESeq2 were left to default.

To assess pathway activity, differentially expressed genes were ranked in decreasing order by their log fold-change compared to baseline. To ascertain whether our fold-changes were within biological plausibility, we repeated differential gene expression using shrunken log-FCs (using DESeq2’s lfcShrink^38^), and again by first eliminating genes whose raw counts totaled 0 across all animals at baseline. Both methods yielded concordant results with our initial analysis.

This ranked list was then input to GSEA Pre-Ranked (version 4.2.3) using a pre-compiled set of pathways as our reference gene set database, and default parameters ^39,40^. To focus on pathways upregulated early post-prime, those with a significant upregulation (i.e. a false discovery rate (FDR) ≤ 0.25) in hours 1,3, and 6, were plotted as a timecourse for each tissue, using ggplot2 (version 3.4.4) in R ^41^.

Further, to determine leading edge genes for each pathway (for each tissue, at each timepoint), we used GSEA’s leading edge tool on our previously computed GSEA outputs ^40^. Then, to resolve early gene expression behaviors, leading edge genes from pathways of interest were plotted in terms of their log fold-changes at one hour post-prime ^41^. Raw data are available in GEO under accession number GSE264344.

### Sample collection for immunologic studies

Blood was collected into RPMI 1640 media (Corning) containing 5mM of EDTA (Life Technologies). Lymphocytes were isolated using Ficoll-Hypaque (GE Healthcare) density centrifugation. The interphase was collected into R10 media, washed, and isolated cells were then used for subsequent assays.

For early timecourse studies muscle and draining lymph nodes were collected into R5 media (RPMI (Corning), 5% FBS (Sigma), 1% Pen/Strep (Life Technologies)). Tissue samples were cut into pieces and placed into R5 media containing collagenase Type IV (Sigma), and then digested for one hour at 37°C on a rocker. Following digestion, samples were passed through a 70 µm filter and any remaining pieces were ground and washed through the filter with R5. All samples were washed once and resuspended in R10 media (RPMI, 10% FBS, 1% Pen/Strep) containing Benzonase (Millipore).

For evaluation of day 60 T cell responses via tetramer staining, collected tissues were harvested and collected into R10 media (RPMI (Corning), 10% FBS (Sigma), 2% pen/strep (Life Technologies). Spleen and draining lymph node samples were ground through 70 µm filters. Spleen samples were treated once with 1X ACK lysis buffer to remove red blood cells. All samples were washed with R10 and passed through a 30 µm filter. Samples were resuspended in R10 media containing Benzonase (Millipore).

### Cytokine and chemokine assays

Frozen serum samples were thawed on ice and subsequently centrifuged for 10 minutes at 10,000 rpm. Serum was treated with 0.05% Tween-20 (Sigma) in 1X DPBS (Life Technologies) for 15 minutes at room temperature. Cytokine and chemokine levels were assessed using the Milliplex Mouse 32-plex premix kit (Millipore) as per manufacturers’ instructions. Samples were subsequently fixed with 2% formaldehyde in 1X DPBS (Life Technologies) for one hour at room temperature. Following, samples were washed, resuspended in Drive Fluid (Luminex Corp.), and run on a Magpix with Xponent software (Luminex Corp). Data was analyzed using a 5-parameter logistic model with an 80-120% standard acceptance range. Extrapolated data below the limit of quantification were graphed and analyzed at the Lower Limit of Quantification for the specific analyte.

### Flow cytometry

Single cell suspensions were first stained with Fixable Blue or Near-IR vital dye in 1X DPS (Life Technologies) for 20 minutes at 4°C. Samples were subsequently washed, blocked with Fc block (TruStain FcX PLUS anti-CD16/CD32, Biolegend) and monocyte block (True-stain monocyte blocker, Biolegend) at 4°C for 15 minutes, then stained with surface antibodies in MACS buffer (MACS wash buffer (Miltenyi Biotec), BSA (Mitlenyi Biotec), Pen/Strep (Life Technologies)) and Brilliant Stain Buffer Plus (BD Biosciences) for 60 minutes at 4°C. For innate profiling experiments antibody panels included: CD45 (clone 30-F11), B220 (RA3-6B2), CD8a (53-6.7), CD80 (16-10A1), CD86 (GL1), Sirpα/CD172a (P84), F4/80 (BM8), CD11c (N418), CD19 (clone 6D5), CD3 (clone 145-2C11), NK1.1 (PK136), CD103 (2E7), MHC II (M5/114.15.2), CD64 (X54-5/7.1), CD40 (3/23), DEC205 (NLDC-145), Langerin (4C7), XCR1 (ZET), Ly6C (AL21), CD11b (M1/70). For tetramer staining experiments antibodies included: CD8a (53-6.7), CD44 (IM7), AL11 tetramer. AL11 monomers were provided by the NIH tetramer core facility (Emory University, Atlanta, GA) and tetramerized using streptavidin conjugated to Brilliant Violet 421 (Biolegend). All antibodies were obtained from Biolegend, or BD Biosciences. Following staining samples were washed and fixed with 2% formaldehyde. Data were acquired on a FACSymphony (BD Biosciences) or LSR II (BD Biosciences) using BD Diva software and analyzed using FlowJo v10 (Treestar).

### Statistics

Statistical analyses on immunologic data were performed using Graphpad Prism 7, using tests as indicated in the text and corrected for multiple comparisons where indicated.

